# Evaluation of bacterial proliferation with a microfluidic-based device: Antibiochip

**DOI:** 10.1101/792358

**Authors:** Valentina Gallo, Alessia Ruiba, Massimo Zanin, Paolo Begnamino, Sabina Ledda, Tiziana Pesce, Giovanni Melioli, Marco Pizzi

## Abstract

The measurement of the proliferation (and the relevant inhibition of proliferation) of microbes is used in different settings, from industry to laboratory medicine. Thus, in this study, the capacity of the Antibiochip (ELTEK spa), a microfluidic-based device, to measure the amount of *E. coli* in certain culture conditions, was evaluated. An Antibiochip is composed of V-shaped microchannels, and the amount of microparticles (such as microbes) is measured by the surface of the pellet after centrifugation. In the present study, different geometries, volumes and times were analyzed. When the best conditions were identified, serial dilutions of microbial cultures were tested to validate the linearity of the results. Then, with the use of wild *E. coli* strains isolated from medical samples, the relationship between bacterial susceptibility to antibiotics (gentamicin, amikacin and ceftriaxone) measured by standard methods and that measured by the Antibiochip was evaluated. In this report, the good quality performances of the methods, their linearity and the capacity to identify susceptible microbial strains after 60 minutes of incubation are shown. These results represent a novel approach for ultrarapid antibiograms in clinics.

## Introduction

During a series of studies focused on the possible uses of micro- and nanochannels in laboratory diagnostics, a group of researchers from ELTEK S.p.A. (Casale Monferrato, Italy) developed and patented a system characterized by an array of microchannels, in which the quantitation of the particulate fraction of a suspension is measured by optical means. The method is based on different V-shaped microchannel arrays (VSMCAs). When microchannels are filled with a solution containing a particulate fraction, after one or more rounds of centrifugation, the volume occupied by the pellet is evaluated with a microscope and an image analyzer. Interestingly, this method is so sensitive and reproducible that small modifications of the fraction of the particulate can be detected. Microbe and yeast amounts represent substrates that are commonly measured both in industry and in medicine. In this report, we describe the first results of bacterial quantitation obtained using wild strains of the human pathogen *Escherichia coli*.

## Materials and Methods

### The microchannel array

The VSMCs were built on a plastic disk (diameter 90 mm) by standard photolithographic techniques. Fig. 1 shows the different geometries used, with a focus on the different angles of the VSMCs. In the present set of experiments, the microchannels had a length of 20 mm and a height of 11, 22 or 36 μm. The geometry of the microchannels was defined starting from the industrial feasibility of low-cost fabrication techniques and the filling properties of the channels. Thus, 50 μm wide channels were identified as a suitable starting point. Fig. 2 shows the loading sector of the microchannels (A), the array of microchannels (B), the microbes floating into the microchannels before centrifugation (C) and the microbial pellet after centrifugation (D).

**Figure 1.**
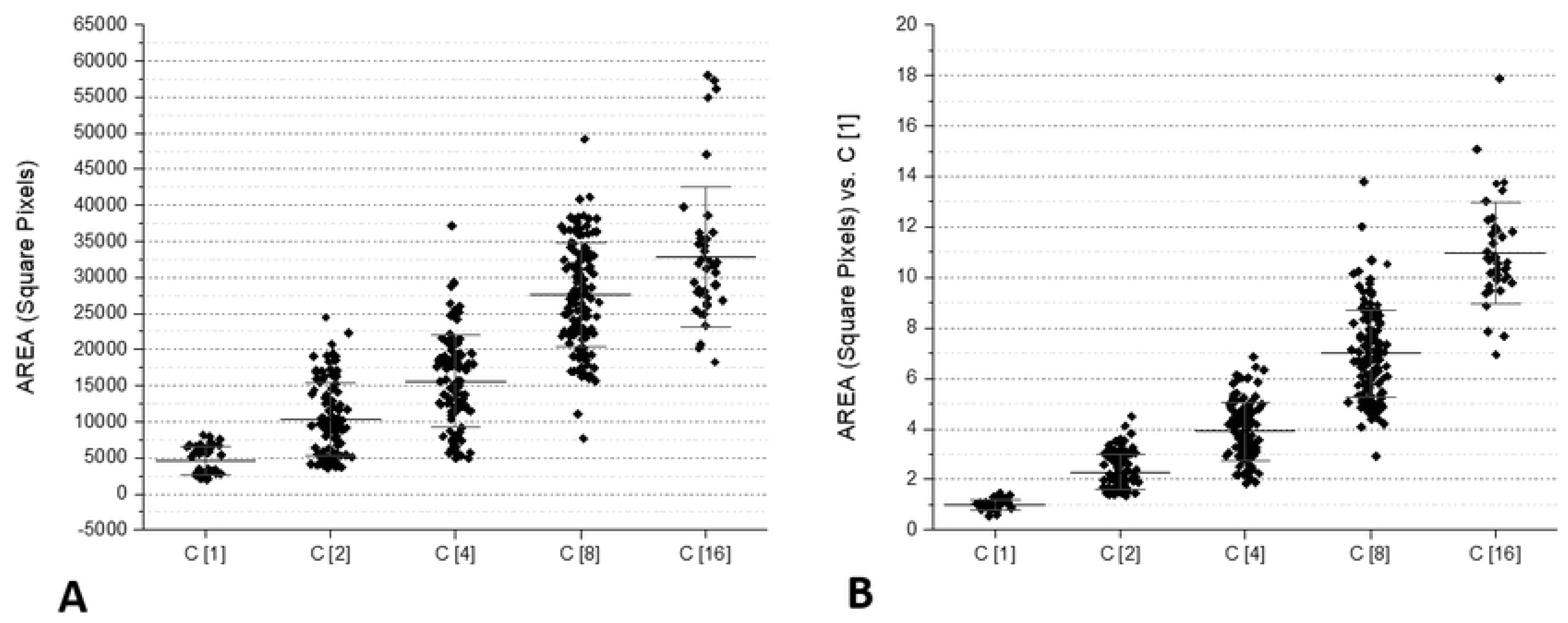
Examples of different geometries and shapes designed and tested during the industrial phase of the development of the VSMCs.

**Figure 2.**
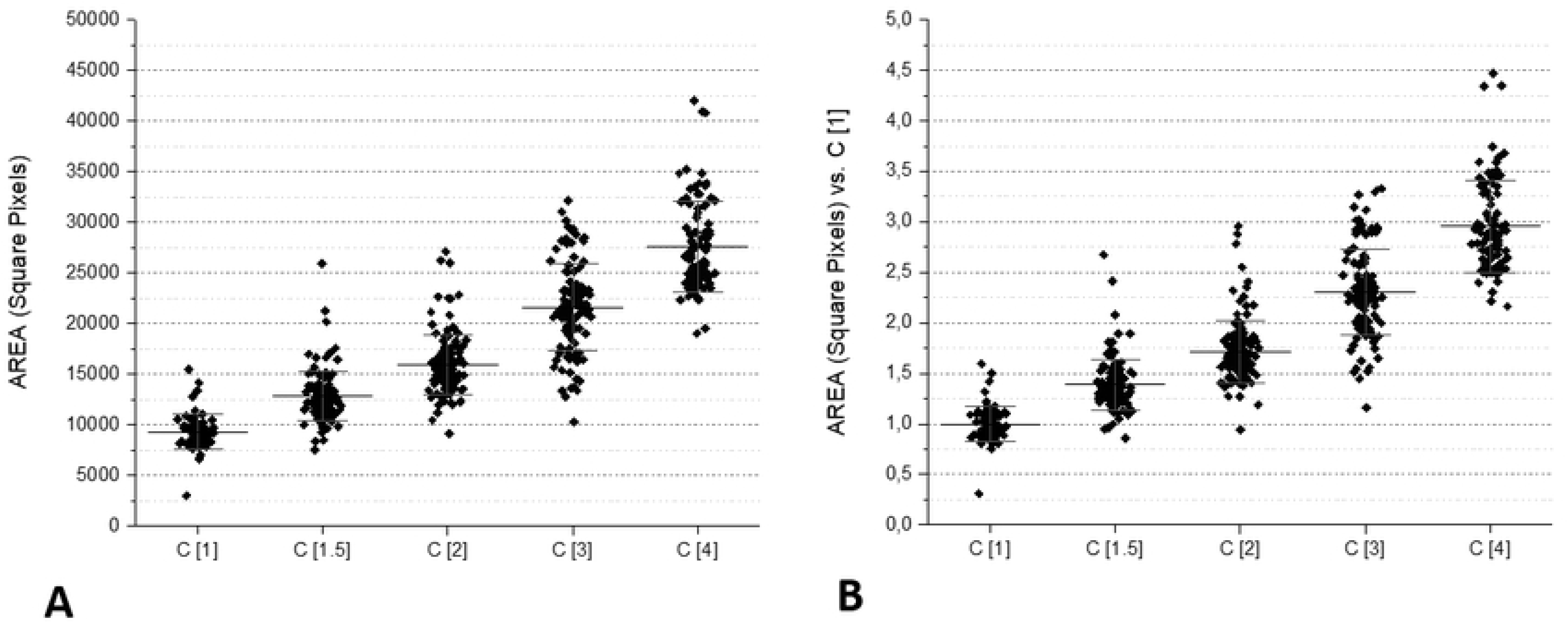
VSMCs at work. (A) Loading sector of microchannels, (B) the array of microchannels, (C) microbes (*E. coli*) floating in the microchannels before centrifugation and (D) the microbial pellet after centrifugation.

To evaluate the quality of the pellets obtained by these different angles and volumes, the following parameters were kept under consideration: 1) the filling time of the channels, important to increase the speed of the test time; 2) the shape of the microbial pellet after centrifugation, important to maintain a regular geometric shape (triangle-like shape) that can be measured by the image-analysis software; and 3) the texture of the microbial pellet after centrifugation, important for the stability of the pellet during the time between centrifugation and the measurement.

### Microorganisms

The microorganisms used in this study were *E. coli* isolated from clinical samples. In detail, the bacterial strains used in this study were obtained from a clinical microbiology laboratory studying samples from hospital and community infections. Colonies were isolated, grown overnight in broth, and then a master microbe bank (MBB) was established and stored at −20°. The bacterial concentration was measured by turbidimetry in McFarland units. Subsequent growth of the bacterial strains was conducted starting from one vial of the MBB. Of all bacterial strains, the identification of the different strains as well as the antibiotic susceptibilities were obtained by a Biomerieux Vitek^®^ 2. Some strains were susceptible to all antibiotics tested, while others had evident resistances.

### Antibiotics

The following antibiotics were used: gentamicin (GEN, Gentalyn MSD vials for human use, 40 mg/mL), amikacin sulphate (AMK, Sigma-Aldrich, cat. A0365900) and ceftriaxone sodium (CTO, Sigma-Aldrich, cat. C5793). All antibiotics were used at the MIC (namely, MIC >4 mg/L for gentamicin, MIC >16 mg/L for amikacin and MIC >2 mg/L for ceftriaxone).

### Measurement of microbial concentration by VSMCs

Microbes were harvested by collecting one or more colonies with an inoculating loop from the agar plates where they were isolated. The colonies were dissolved in 0.5% NaCl, and the bacterial concentration was measured in McFarland units with a Biomerieux densitometer. The bacterial suspension was used when the McFarland number was 1 or more.

In a first series of tests, two-fold dilutions of each microbe were loaded into the microchannel prechamber, and when the microchannels were filled with the suspension, disks containing the VSMCs were centrifuged at 7000 rpm for 3 minutes, according to the operating conditions identified in the preclinical (industrial feasibility) phase of the study. During this set of experiments, different microbes were plated in VSMCs at different angles and volumes.

To detect the presence of the microbial pellet and to calculate the surface of the pellets, a USB-driven 800X magnification objective (AM7515MT8A Dino-Lite Edge Microscope, from AnMo Electronics Corporation, Hsinchu, Taiwan) was assembled in the centrifuge to measure the bacterial pellets. A 3D electric servomechanism was used to align and focus the lens on the VSMCA. The surface of the microchannel occupied by the microbial pellet was measured on a different picture by ImageJ, a free NIH software for image analysis.

Another series of tests was carried out to evaluate whether VSMCs were suitable to detect not only the differences in microorganism concentrations obtained by a serial dilution but also the inhibition of bacterial proliferation caused by antibiotics in susceptible strains. For this, different microbes were incubated in a plastic tube at 37°C in a moist chamber in the presence of the MIC of different antibiotics. After 30 and 60 minutes, a microsample of the culture was collected by a micropipette and loaded into the loading chamber of the VSMC. When the VSMCs were filled, the disks were centrifuged, and the pellets were detected and measured as described above. In this series of tests, both *E. coli* strains susceptible to GEN, AMK and CTO and strains resistant to GEN and CTO (according to standard medical diagnostics procedures) were used.

## Results

Different geometries and volumes were used during the validation phase of the VSMC to define the best configuration of the microchannels. Table 1 shows the results of these tests. The thinnest channels (11-μm-thick) required a long filling time and did not guarantee a proper concentration in the channels when compared with the 22- and 36-μm-thick channels. Similarly, 70° angles were the best, while 90° pellets were unstable and 60° pellets displayed several bubbles in the channels.

**Table 1.** Evaluation of geometries and volumes to define the best configuration of microchannels.

| Parameters evaluated | Thickness of the microchannel<br>(in $\mu\text{m}$) | | | Angle of the V-shaped channels<br>(in degrees) | | |
| --- | --- | --- | --- | --- | --- | --- |
|  | 11 | 22 | 36 | 90° | 70° | 60° |
| Filling of the channels | - | + | ++ | + | + | - (2) |
| The shape of the pellet after centrifugation | +/- | ++ | + (1) | + (1) | ++ | - (2) |
| The texture of the pellet after centrifugation | +/- | + | +/- | + (1) | ++ | - (2) |

(1) Instability of the pellet
(2) Presence of air bubbles in the microchannel.

Due to a different diffusion behavior, even if the size of the channels was approximately 10 times the size of the particles, the concentration inside the thinnest channels did not reproduce the concentration in the sample, as shown in Fig. 3A. Comparing the absolute values of the pellet areas, the behavior of the 22- and 36-μm-thick channels was very similar, while the 11-μm-thick channels showed a lower particle concentration. This is a well-known phenomenon due to anomalous diffusion when the channel size and particle size are not very different [1, 2].

**Figure 3.**
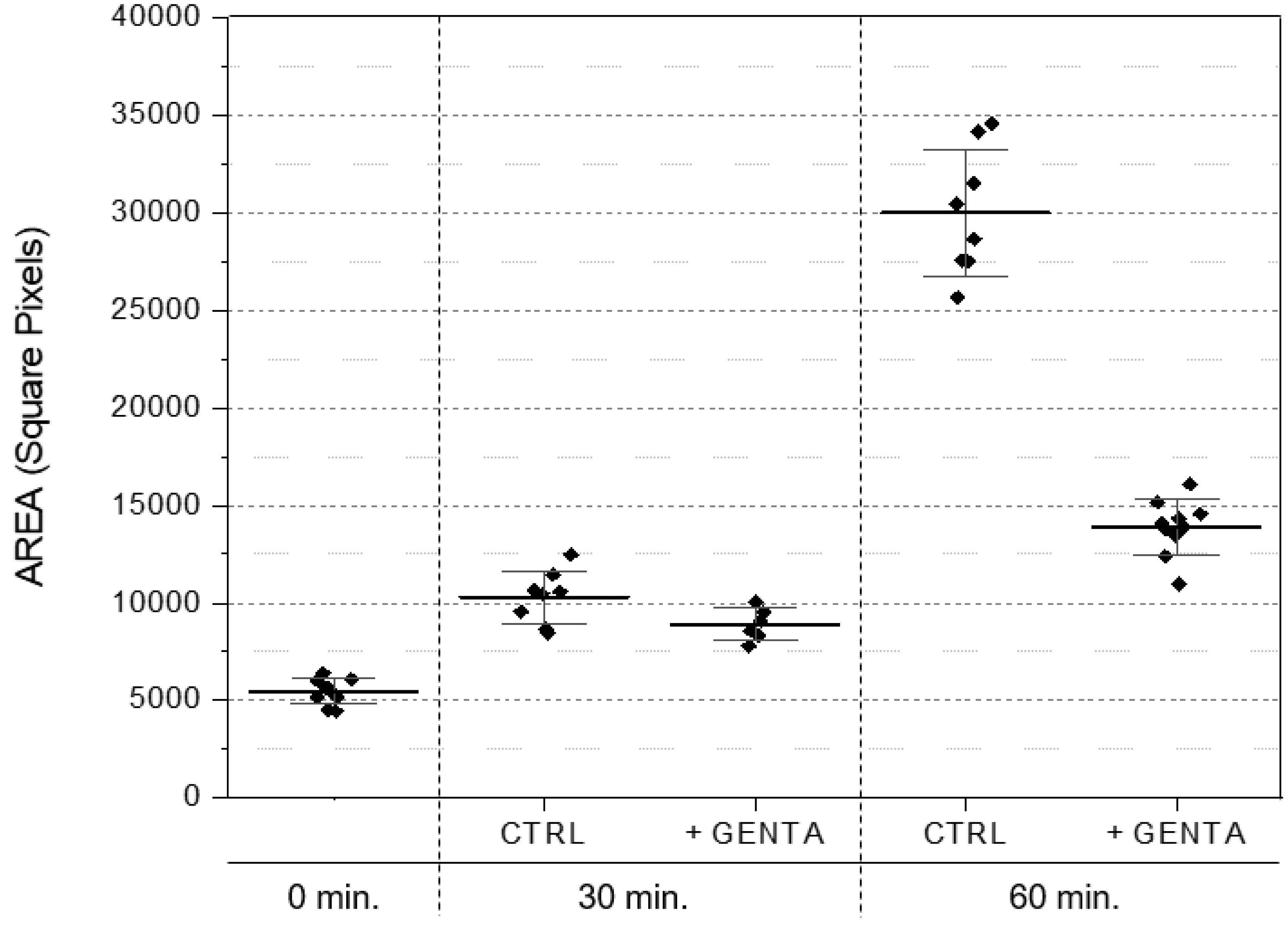
Diffusion of microbes in VSMCs with different thicknesses: (3A) 11μm, (3B) 12μm, (3C) 36μm. Measurement of the areas in three different microbial concentrations (1:4 dilutions). See also the text.

The concentration measurement with 22- and 36-μm-thick channels (Fig. 3B and 3C) was comparable, but the pellet in the 36-μm-thick channels was unstable, leading to a larger area distribution at higher concentrations.

The correlation between the expected and measured relative concentrations of the samples was correct for the 22- and 36-μm-thick channels, while it was not respected for the 11-μm-thick channels, as shown in Fig. 4.

**Figure 4.**
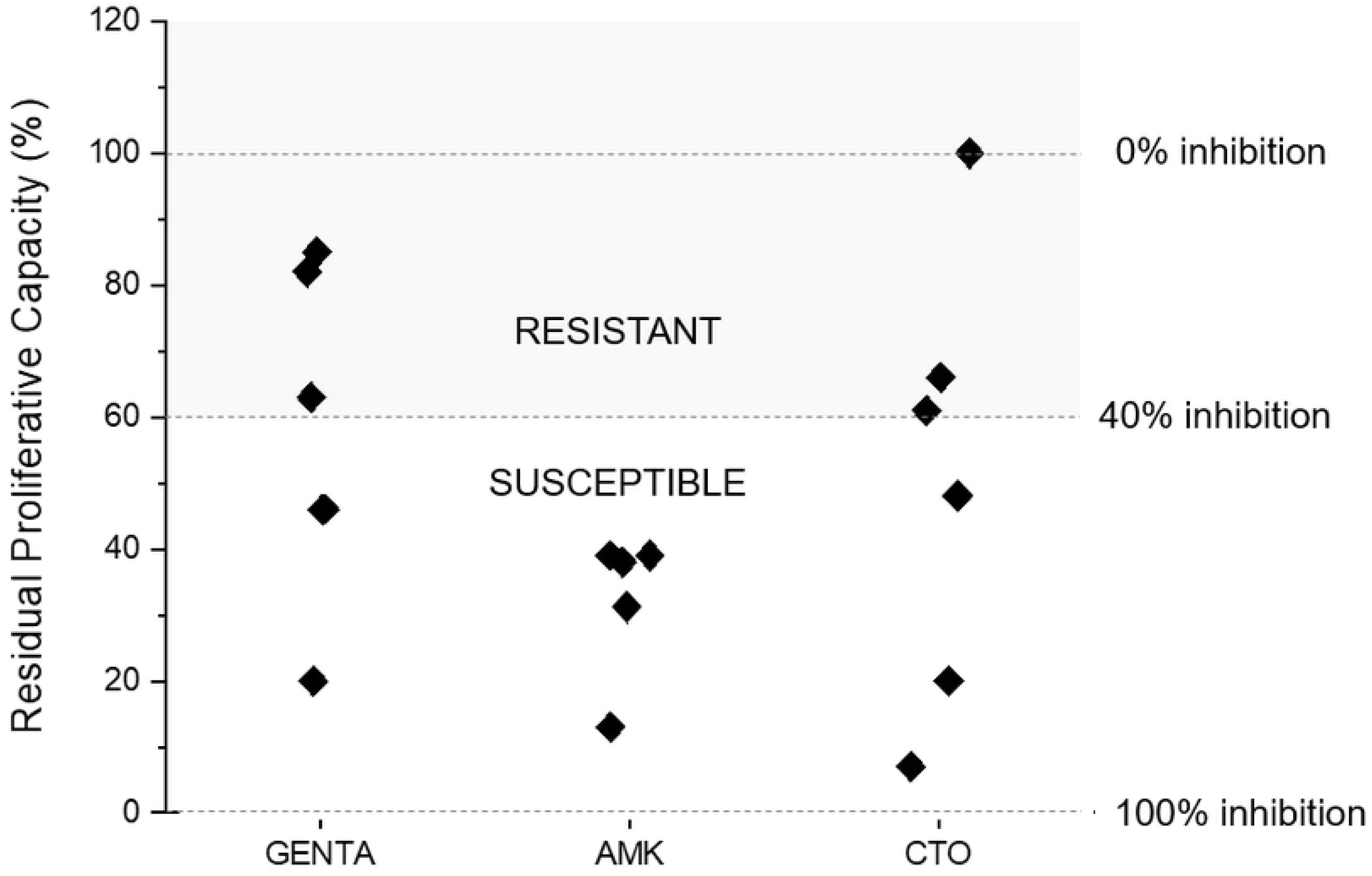
Diffusion of microbes in VSMCs with different thicknesses: (3A) 11μm, (3B) 12μm, (3C) 36μm. Correlation of the areas vs expected results in three different microbial concentrations (1:4 dilutions). See also the text.

Based on these results, subsequent experiments were conducted with the 22-μm-thick VSMC at a 70° angle.

### Dilutions

Several tests were performed using two-fold serial dilutions of a bacterial suspension starting from an optical density > 1 McFarland. Fig. 5 shows the results of these assays. It is evident that the surface of the pellets was in strict correlation with the dilutions of microorganisms. In practical terms, these dilutions corresponded to four replications of the original bacterial suspension; considering that *E. coli* doubles in 20-30 minutes, a 16-fold increase is representative of a culture time from 1.2 to 2 hours. To further explore the capacity of VSMCs to detect a small difference in the microorganism concentration, another series of experiments was carried out using a narrower dilution range, i.e., 1, 1.5, 2, 3 and 4. Fig. 6 shows the results of these tests. In these conditions, it was evident that the surface of the pellets in the VSMC was highly representative of the small differences of the original microorganism suspensions.

**Figure 5.**
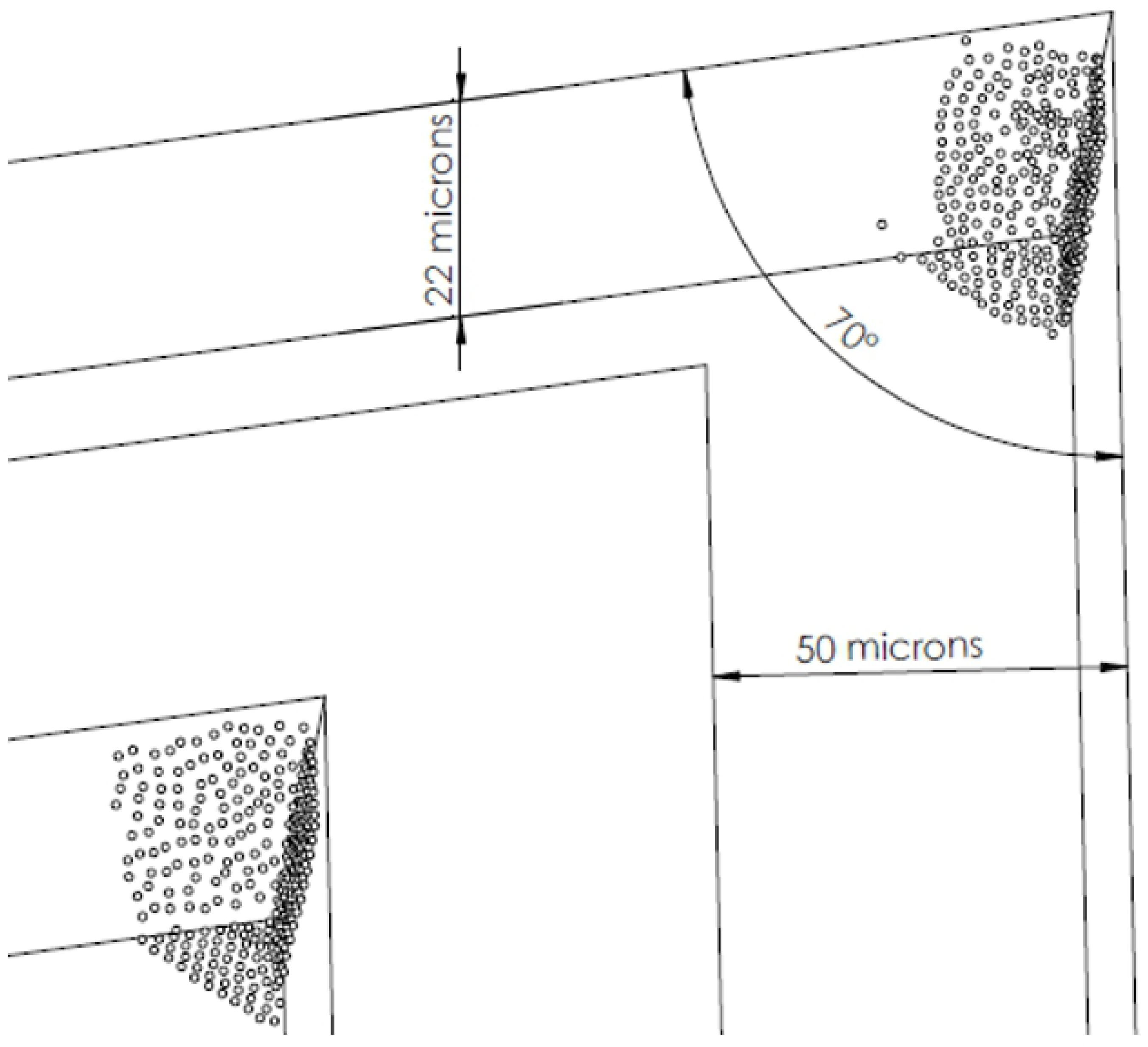
Microbe areas in VSMCs observed with two-fold serial dilutions of a bacterial suspension.

**Figure 6.**
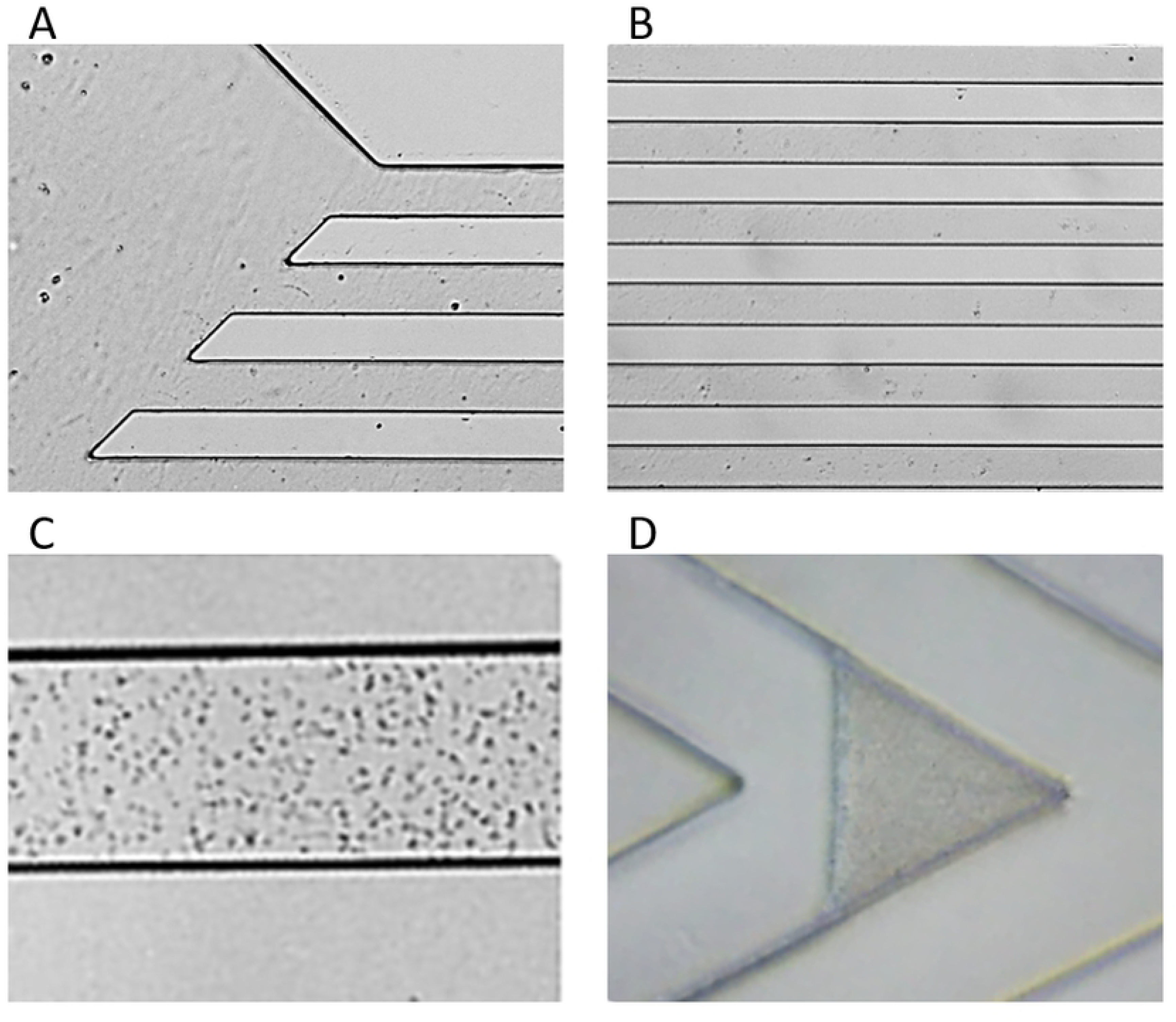
Microbe areas in VSMCs loaded with a narrow dilution range (1, 0.75, 0.50, 0.25, 0.1). In these experiments, different McFarland concentrations (from 1 to 2) were used and normalized for dilutions.

### Inhibition of bacterial proliferation by antibiotics

Starting from the abovementioned results and the clear capacity of VSMCs to detect small differences in the concentration of microorganisms in a suspension, several tests were planned to verify whether the inhibition of proliferation due to one or more antibiotics could be measured when different antibiotics were introduced to susceptible microorganisms. For this, different wild strains of *E. coli*, obtained from clinical samples and fully characterized by standard microbiology methods, were incubated with different antibiotics (gentamicin, amikacin and ceftriaxone) at the respective MIC values. Fig. 7 shows an example of the experiments performed using bacterial strains that were susceptible to antibiotics in routine antibiograms. Once clear evidence of the capacity of VSMCs to detect the inhibition of the proliferation of *E. coli* was observed within 60 minutes of culture, two other wild-type bacterial strains, characterized by resistance to antibiotics (namely, gentamicin and ceftriaxone), were tested and, as expected, different results were observed. Fig. 8 shows a summary of all the results.

**Figure 7.**
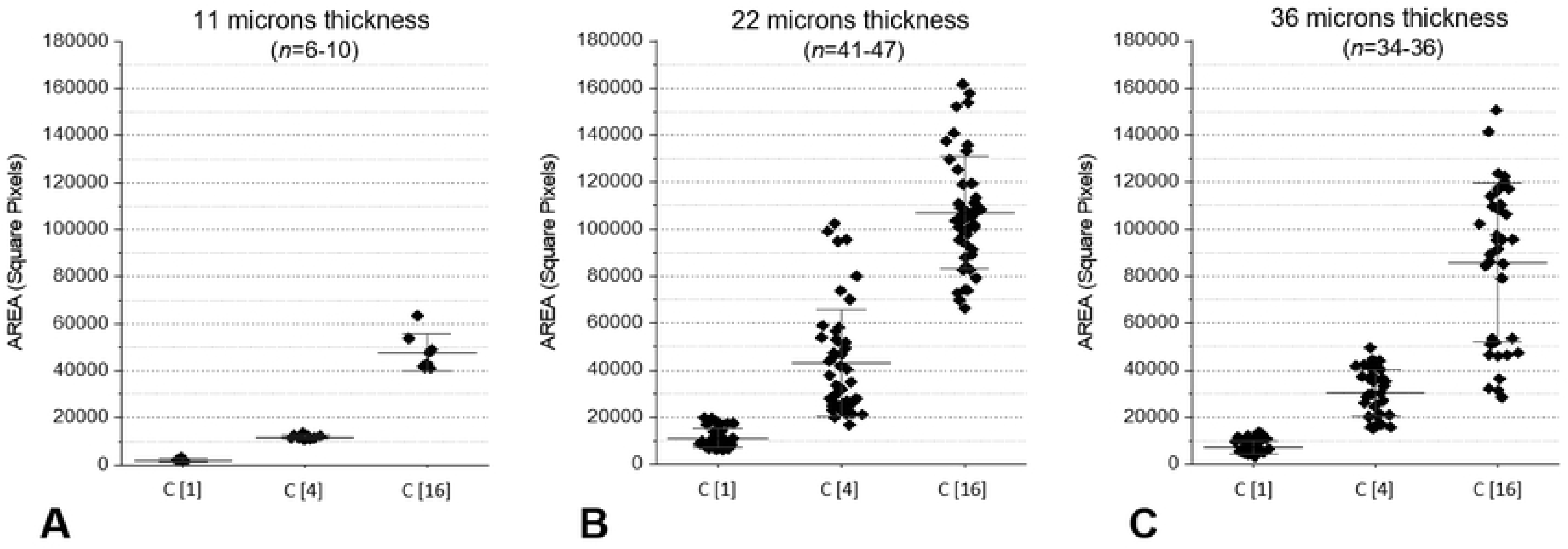
Inhibition of bacterial proliferation in different strains of *E. coli* susceptible to antibiotics. The results were obtained after 30 and 60 minutes of incubation. Susceptible strains were barely evident after 30 minutes but clearly evident after 60 minutes.

**Figure 8.**
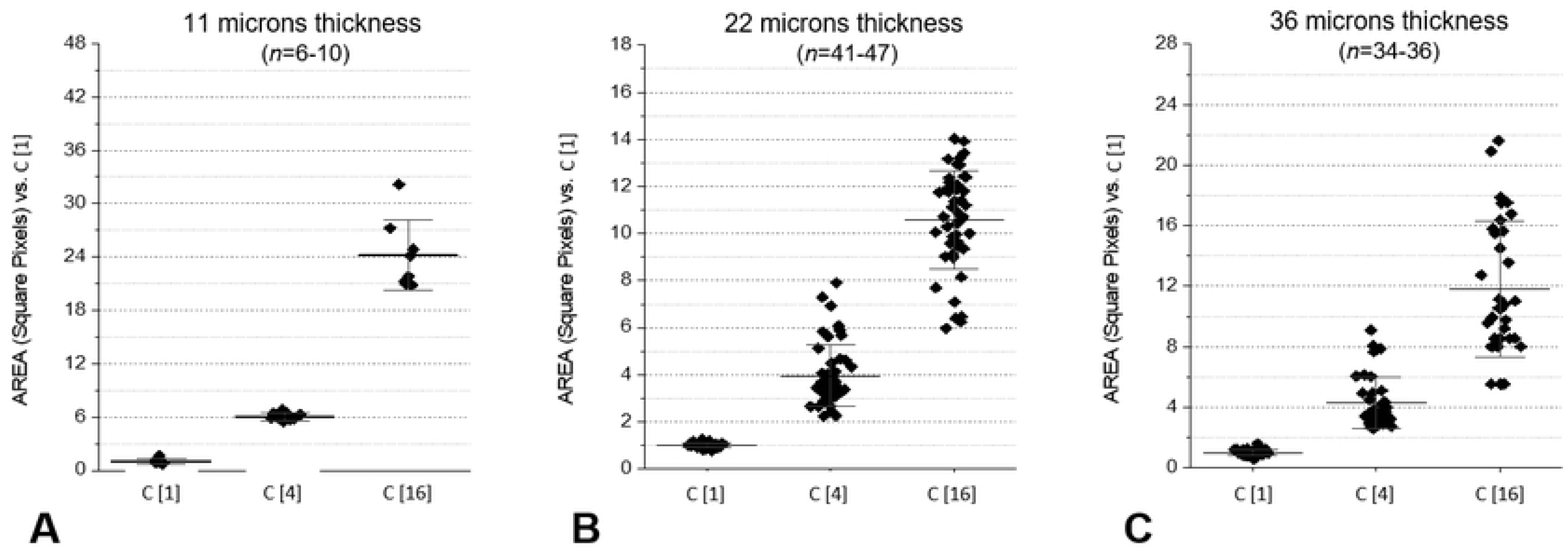
Analysis of *E. coli* strains cultured in the presence of antibiotics; 3/5 were resistant to GEN, all were susceptible to AMK and 3/6 were resistant to CTO in standard microbiology tests. VSMC results correlate well with standard results.

In an attempt to find some general rule, the results of the VSMCs for different antibiotics were grouped according to the antibiotic resistance as detected by the standard method. A simple statistic (mean and standard deviation after transforming the percentages to arcsin) was calculated. The mean result for strains evaluated as susceptible by standard methods (independently from the antibiotic used) was represented by a capacity of proliferating of 25% (corresponding to a 75% inhibition), with an upper limit of 60% (40% inhibition). For resistant strains (at the MIC), the proliferating capacity was 85% (15% inhibition) with a lower limit of 60%. According to these results, the presence of a proliferative capacity < 60% was considered to correspond with the susceptibility to the standard tests, while the maintenance of a proliferative capacity > 60% of the controls was considered suggestive of antibiotic resistance.

While the large majority of experiments were carried out using *E. coli* (mainly wild-type), other experiments were conducted with other gram-negative bacteria, such as *Enterobacter aerogenes, Klebsiella pneumoniae* and *Proteus mirabilis* (not shown). At present, the number of these tests is too small to attempt any statistical consideration, but from a descriptive point of view, these other Bacteriaceae also had a behavior comparable to that of *E. coli* in the VSMC model. In the same series of experiments, other incubation times were also considered, in particular 30 minutes and 120 minutes. The 120-minute incubation time confirmed the results observed at 60 minutes, and for this reason, at least in the present setting of assays, no further tests were carried out. In contrast, highly promising results were observed after 30 minutes of culture in the presence of antibiotics; nevertheless, the number of tests performed does not seem to be sufficient to provide some shareable conclusions.

## Discussion

Methods to measure the capacity of an antibiotic to inhibit the proliferation of a bacterial strain have been developed in the past, starting from the first description by Fleming [3] to the antibiotic diffusion in paper disks [4] and the more recent measurement of the minimal inhibitory concentration [5]. The recently introduced technology by Accelerate Diagnostics uses morphokinetic cellular analysis to measure the capacity of a microbe to proliferate in the presence of an antibiotic [6]. At present, there is evidence that increasing the speed of the detection of antibiotic resistance is a real topic [7], [8, 9], [10], [11].

In routine clinical microbiology settings, the time to measure antibiograms from an automated or manual system ranges from 6 to 12 hours. In some cases, for example, with the use of Accelerate or by the use of the Alfred 60AST (Alifax, Nimis, IT) culture system, results can be observed in a shorter time. Of note, this “laboratory time” is added to the period (usually, one night) used to isolate a microbe from a biological sample. The rapid methods currently available also require a specific laboratory organization possible only in highly specialized structures. Nevertheless, patients with infectious diseases or patients with severe systemic diseases complicated with a concomitant infection require a very rapid description of the susceptibility of the microbe to start a specific antibiotic therapy in a very short time. This medical emergency is complicated by the diffusion of antibiotic resistance [12] and by the fact that new antibiotics do not seem to be in the industrial pipelines [13].

The VSMC method described above seems to be different from others and highly innovative. Indeed, the tests described in this work show that the surface of the pellet, strictly related to the number of microbes, is detected rapidly and accurately. The reproducibility is good, and the correlation between measured surfaces and expected results – at least based on two-fold dilution assays – is excellent. In addition, other more accurate measurements (performed using a 25% increase in the bacterial concentrations) also showed that the capacity to distinguish between apparently small differences is good. This seems to be even more relevant considering that the prototypic gram-negative microbe *E. coli* doubles in 20-30 minutes. Thus, the increase in the number of microbes – paralleled by the increase in the surface of the pellets – ranges between 2- to 3-fold in a period of one hour, and an increase in the number of microbes measured using VSMC can be detected in 30 minutes. Notably, these results can be observed by starting the microbial culture with a concertation of microbes ranging from 0.5 to 1 McFarland, a routine concentration of microbes for automated clinical tests, easily achievable by harvesting more than one colony from the isolation plate.

By the use of VSMCs, microbes susceptible to antibiotics, cultured with and without antibiotics, had different pellet areas in VSMCs after 60 minutes, and some preliminary data indicate that 30 minutes may be enough time to observe this effect. The microbial proliferation in negative (no-ATB) control tubes is regular and easily detectable by the measurement of the VSMC. The same microbes, in the presence of an antibiotic, are inhibited, and significant modifications of the pellet surfaces can be measured. Therefore, the method based on VSMC, herein described, seems to be able to identify antibiotic-susceptible microbes in laboratory tests lasting a period of time (namely, 60 minutes) significantly different from other routine tests used in clinical microbiology. In an attempt to compare the results of the two approaches, an extremely simple statistical approach was used, and a clear correlation between antibiotic susceptibility was observed. As soon as many other results are obtained using different microbes and different antibiotics, a much more sophisticated statistical analysis will be performed. However, although it is probable that the percentages of proliferation will be more accurate, it seems that deeply modifying the general concept will be difficult.

Unless there is a wide understanding of the evaluation of fine modifications of microbial inhibition in short culture times, VSMC cannot be used in the present form to measure the MIC of single antibiotics. However, the standardization of the culture conditions (namely, an accurate microbial concentration for each different strain, optimized culture and centrifugation conditions, and a highly automated and fast reading system) would, in the future, allow the use of VSMC for the direct detection of MIC values (or, as a second possibility, calculation) as made currently by the methods approved by national regulatory agencies. In medical microbiology, antibiograms are performed not only to identify resistances but also to identify susceptibility to antibiotics. Indeed, a basic medical rule is to administer patients with an antibiotic that may have a high probability of being efficient in controlling the infection. Resistances are extremely relevant from an epidemiological point of view, but in the real time management of the patient, they seem less relevant.

In the present experimental phase of analysis of microbial proliferation (and inhibition) in VSMCs, specific warnings must be given. First, for microbes that were resistant to standard clinical methods, a proliferation identical to that observed for the controls was never observed. In other words, even for “resistant strains”, a certain susceptibility is detected in short time tests. Second, antibiotic susceptibility can be identified with great accuracy, even if a certain residual proliferative capacity is detected in 60 minutes of incubation. In these conditions, the VSMC-based method detects something that is invisible in the long-term culture (>6 hours) used by the standard methods, where antibiotics have plenty of time to kill all microbes.

Third, antibiotic resistance seems more difficult to define in very short-period assays. Indeed, a certain inhibition is always detected, even if a correlation between certain behavior patterns of “resistant” microbes measured by VSMC and “classic” routine tests was observed. For example, the introduction of a cut-off of 30% (i.e., resistant microbes in the presence of antibiotics maintained at > 70% capacity of proliferation when compared with the “negative control”) indicates that even at an MIC of 1, a certain inhibition of resistant microbes is observed in the short test. This is in partial contrast with the resistant/susceptible paradigm of clinical microbiology. In the same context, one may argue that one of the postulates of clinical microbiology is that a single resistant microbe in a colony of susceptible microbes means that the whole colony should be considered resistant. This can be easily observed in long-term (such as overnight) cultures of microbes, where it is virtually impossible to detect the fraction of resistant microbes in a colony, even if a single mutation is possible but not very frequent. When short time tests are used, the rapidity of the results to the clinician is counterbalanced by the objective difficulty to identify extremely rare microbes carrying a resistance phenotype in the context of a colony. Clearly, this is not a specific problem of the method described in this work, but it is in common with all other attempts to increase the speed of antibiograms. However, even if the problem is considered from a medical point of view, it may be expected that the very large majority of susceptible microbes are kept under control by the antibiotic treatment and, probably, the remaining resistant bacteria can be controlled by an immediate immune-response that has a relevant role in managing bacterial infections. Nevertheless, the presence of a selective agent (the antibiotic) administered to the patient cannot rule out that resistant microbes may proliferate.

In conclusion, many attempts have been made to reduce the time of incubation of microbes and to detect microbial susceptibility or resistance to antibiotics. In the present study, very short incubations (60 minutes), unimaginable a few years ago, have been shown to be possible to detect microbial susceptibility or resistance to antibiotics, provided that an extremely sensitive method of bacterial proliferation/inhibition is used. However, the clinical impact of this new technology in the control of infectious disease must be carefully evaluated. Obviously, if the described correlation between the VSMC method and classic methods is confirmed, clinical trials will be much safer to be carried out in complex patients with severe infections to evaluate whether a shorter analytical time has an impact on the patient’s clinical evolution.

